# Antisense oligonucleotide therapy for the common Stargardt disease type 1-causing variant in *ABCA4*

**DOI:** 10.1101/2022.08.12.503728

**Authors:** Melita Kaltak, Petra de Bruijn, Davide Piccolo, Sang-Eun Lee, Kalyan Dulla, Thomas Hoogenboezem, Wouter Beumer, Andrew R. Webster, Rob W.J. Collin, Michael E. Cheetham, Gerard Platenburg, Jim Swildens

## Abstract

The c.5461-10T>C p.[Thr1821Aspfs*6,Thr1821Valfs*13] variant has been identified as the most common severe Stargardt disease type 1 (STGD1)-associated variant in *ABCA4*. STGD1 is the most recurrent hereditary form of maculopathy and so far, no treatment is available for STGD1. In STGD1 patients homozygous for this variant, the onset of the disease typically is in childhood and patients are legally blind by early adulthood. The variant leads to exon skipping and generates out-of-frame *ABCA4* transcripts that prevent the translation of functional ABCA4 protein.

We applied antisense oligonucleotides (AONs) to restore the wild-type RNA splicing in *ABCA4* c.5461-10T>C. The effect of AONs was investigated *in vitro* using an *ABCA4* midigene model and 3D human retinal organoids (ROs) homozygous for the *ABCA4* c.5461-10T>C variant. The mRNA in untreated ROs contained only disease-associated isoforms, whereas the organoids treated with the lead AON sequence showed 53% splicing correction and restoration of ABCA4 protein.

Collectively, these data identified the lead candidate QR-1011 as a potent splice-correcting AON to be further developed as therapeutic intervention for patients harboring the severe *ABCA4* c.5461-10T>C variant.

## INTRODUCTION

Clearance of toxic retinoid metabolites from the visual cycle is essential for the maintenance of functional retinal pigment epithelium (RPE) and the underlying cone and rod photoreceptor cells in the retina. This is in part carried out by the ATP-binding cassette, sub-family A, member 4 (ABCA4) protein, which is localized in the outer segment disk rims of photoreceptor cells. In absence of functional ABCA4, N-retylidene-PE and the lipofuscin fluorophore A2E accumulate in the RPE, leading to the death of the RPE layer and the photoreceptor cells (Allikmets et al. 1997; Mata, Weng, and Travis 2000; Quazi, Lenevich, and Molday 2012; Weng et al. 1999). In the case of two *ABCA4* null-alleles, disease progression is rapid and clinically recognized as autosomal recessive cone-rod dystrophy (CRD), whereas the involvement of alleles with residual ABCA4 function gives rise to Stargardt disease type 1 (STGD1) (Cremers et al. 1998; Maugeri et al. 1999; Sangermano et al. 2016). Although STGD1 is the most common form of inherited macular dystrophy, no treatment options are available, highlighting the importance of the development of therapeutic strategies. Studies on large cohorts of STGD1 patients have identified more than 2,200 disease-associated variants in *ABCA4* (Cornelis et al. 2022), the majority of which are missense variants, followed by mutations that alter pre-mRNA splicing (Cornelis et al. 2017; Paloma et al. 2001; Jonsson et al. 2013; Klevering et al. 2004).

*ABCA4* c.5461-10T>C p.[Thr1821Aspfs*6,Thr1821Valfs*13] is a non-canonical splice site variant that causes the skipping of either exon 39 or exons 39 and 40 together, resulting in the production of out-of-frame *ABCA4* isoforms (Aukrust et al. 2017; Maugeri et al. 1999; Sangermano et al. 2016). It is the most common severe STGD1-causing variant (Cornelis et al. 2022; Kitiratschky et al. 2008; Miraldi Utz et al. 2014; Runhart et al. 2021). The onset of the first symptoms of STGD1 disease in individuals homozygous for the *ABCA4* c.5461-10T>C variant is often in the first decade of life, after which the progress of the disease can sharply accelerate and lead to the state of legal blindness between the age of 20 and 30 (Cideciyan et al. 2009; Sangermano et al. 2016). The steep deterioration of visual acuity in homozygous individuals is explained by dramatically reduced levels of wild-type ABCA4 protein (Aukrust et al. 2017; Huang et al. 2021; Sangermano et al. 2016).

The eye is an isolated and, consequently, immune-privileged organ, which makes it an attractive target for the development of genetic therapies; in fact, the feasibility of gene replacement therapy has been shown by voretigene nepavovec, an FDA-approved treatment for inherited retinal dystrophy (IRD) caused by autosomal recessive variants in *RPE65* (Maguire et al. 2019). However, considering the size of its coding sequence, *ABCA4* remains a challenge for introduction via adeno-associated vectors, which are typically used for gene augmentation therapy. Since many STGD1 disease-causing variants are known to hamper the splicing process and give rise to in-frame or out-of-frame truncations or pseudo-exon insertions in *ABCA4* mRNA, antisense oligonucleotides (AON) are a promising therapeutic strategy due to their ability to manipulate the aberrant splicing and increase the production of functional protein.

AON-induced splice modulating activity showed promising therapeutic strategies by exon exclusion, pseudo-exon exclusion and allele-specific degradation of aberrant transcripts for several IRDs, such as autosomal recessive Leber congenital amaurosis, autosomal recessive Usher syndrome type 2A, inherited optic neuropathy and autosomal dominant retinitis pigmentosa (Adamson et al. 2017; Bonifert et al. 2016; Collin et al. 2012; Gerard et al. 2012; Murray et al. 2015; Slijkerman et al. 2016; Dulla et al. 2018; Parfitt et al. 2016; Russell et al. 2022). The ABCA4 protein consists of two transmembrane domains that harbor 6 transmembrane helices each; any truncation within these sites would severely disrupt the complex protein conformation that is required for its correct function (Figure 1A) (Xie et al. 2021). Hence, to alleviate the STGD1 phenotype, AON-based intervention would need to redirect the reading frame to its original phase. Previous research showed AON-mediated correction of splicing defects in *ABCA4* for several deep-intronic disease-causing variants in midigene-models, differentiated photoreceptor progenitor cells (PPCs) and retinal organoids (ROs) (Bauwens et al. 2019; Khan et al. 2020; Sangermano et al. 2019). To rescue the splicing defect caused by c.5461-10T>C, we designed AONs that exert their action through the mechanism of re-inclusion of skipped exons in *ABCA4* (Figure 1B). This AON-guided mechanism was successfully applied with Nusinersen, the first drug approved for the treatment of spinal muscular atrophy (SMA). Nusinersen is administered intrathecally to reach the cerebrospinal fluid and it targets the intronic splice silencer to re-introduce *SMN2* exon 7 (Hache et al. 2016). The use of this splicing manipulation for the treatment of retinal diseases has not yet been reported.

**Figure 1.**
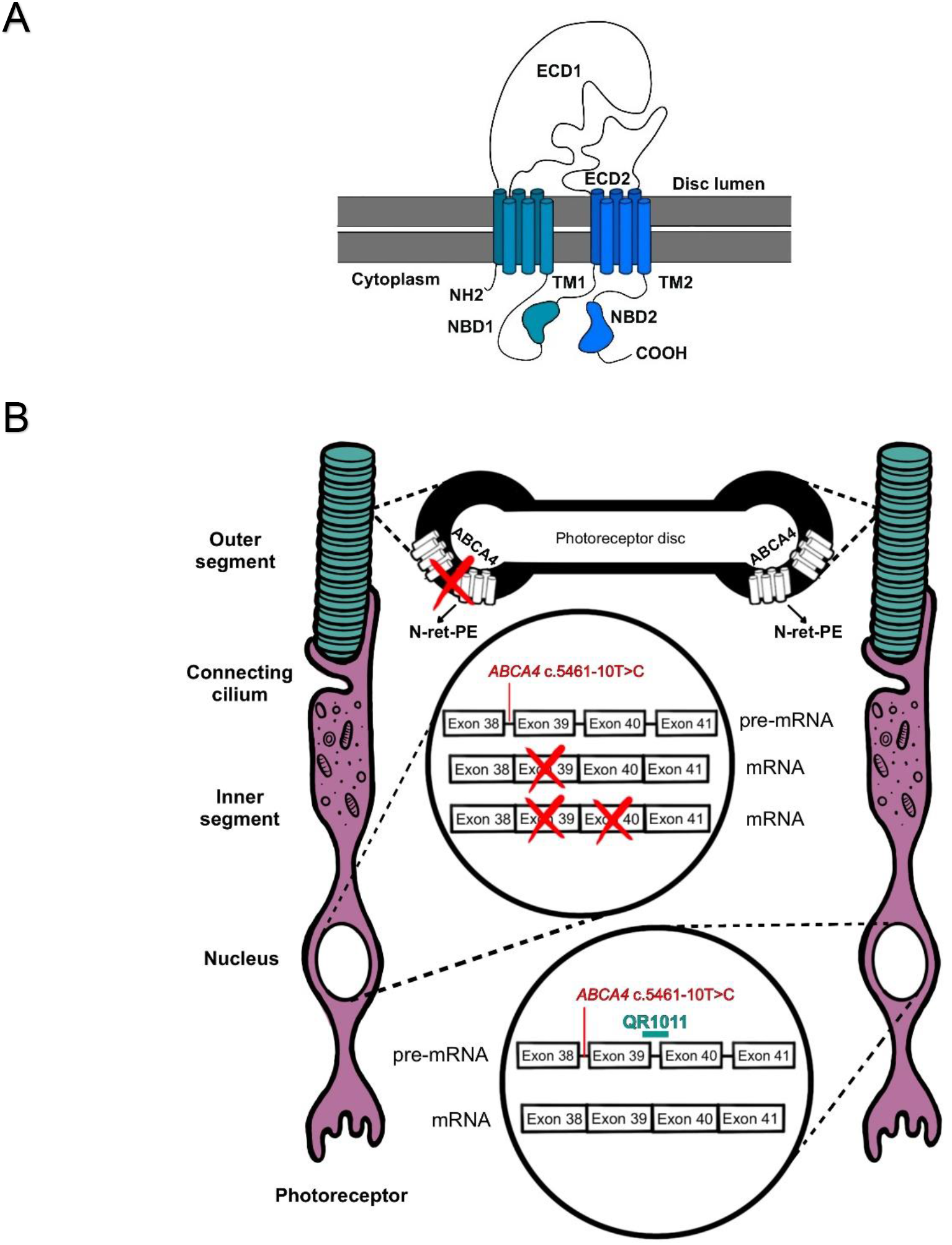
Schematic representation of ABCA4 and the splicing modulating effect of QR-1011. (A) The ABCA4 protein is composed of 2,273 amino acids, and its complex structure involves two transmembrane domains (TM1 and TM2), each with 6 transmembrane helices. In addition, the protein structure displays 2 glycosylated extra cytoplasmatic domains (ECD1 and ECD2) that extend 120 Å from the disk rim [48] and 2 nucleotide binding domains (NBD1 and NBD2) where ATP hydrolysis takes place. Molar ratio of ABCA4 to rhodopsin is present at 1:300 in mouse rod outer segments [47]. (B) In presence of the frameshift variant *ABCA4* c.5461-10T>C, generated transcripts lack either the exon 39 or exons 39 and 40; this splicing defect hampers the production of functional ABCA4 protein and toxic retinoid products (N-retylidene-PE) cannot be removed from photoreceptor’s outer segments, leading to accumulation of A2E and lipofuscin granules, key pathogenic features for STGD1. The splicing modulating the activity of QR-1011 is designed to restore wild-type splicing and include exons 39 and 40 in *ABCA4*. In this way, the functionality of wild-type protein is restored.

Considering the lack of animal and *in vitro* models endogenously expressing *ABCA4* c.5461-10T>C for screening therapeutic molecules, we first assessed splicing in a midigene model incorporating the *ABCA4* genomic region of interest (Sangermano et al. 2018). To assess the effect of AON treatment on ABCA4 protein expression, we employed ROs differentiated from CRISPR-Cas9-edited or patient-derived human induced pluripotent stem cells (iPSCs) (Hallam et al. 2018; Nakano et al. 2012). ROs have been shown to be robust models to investigate possible therapies for various IRDs, considering that they express the targeted variants in the wider genomic environment and their retina-like lamination allows insight into the trafficking and function of disease-associated protein variants (Parfitt et al. 2016; Nakano et al. 2012; Zhong et al. 2014). Additionally, several transcriptomic analyses confirmed high similarities in their key characteristics with native fetal and adult human retina (Kaya et al. 2019; Kaewkhaw et al. 2015; Kim et al. 2019), which, all together, suggest the possibility of ROs replacing animal models to validate the efficacy of retina-targeted therapeutic interventions. Here, we report the ability of an AON to restore c.5461-10T>C *ABCA4* exon inclusion and ABCA4 protein in an RO model of this common variant.

## RESULTS

### Selection of lead AON candidates in *ABCA4* minigene and midigene model

We set out to identify AONs capable of correcting the splicing defect caused by *ABCA4* c.5461-10T>C. Therefore, 31 AONs were designed as an oligo-walk covering exon 39 and the surrounding intronic sequences. During the sequence design, we kept optimal AON parameters when it was possible, such as a GC content of >40% and the Tm >48°C, as described in Aartsma-Rus (2012), to avoid potential off-targets. *ABCA4* is retina-specific, so in absence of a cell line expressing the *ABCA4* c.5461-10T>C variant, the splice-modulating effect of the AONs was assessed in a minigene carrying the *ABCA4* exon 39 and parts of the flanking introns with c.5461-10T>C (Figure S1A), while the construct was flanked by rhodopsin (*RHO*) exon 3 and exon 5 (Sangermano et al. 2018). The minigene was expressed in HEK293 cells that do not express detectable levels of *ABCA4* transcript endogenously. These cells were transfected with 100 nM AON and after 48 hours, transcripts were quantified by digital droplet PCR (ddPCR). The treatment revealed an increase of exon 39 containing transcripts with AONs targeting a region at the 5’ end of intron 39 (Figure S1B) that contains strong intronic splicing silencer motifs (Figure S1C). In particular, AONs 31 and 32 managed to restore 27±1.5% and 21±1% of the *ABCA4* exon 39 containing transcript and served as a basis for further optimization. Since the minigene model did not contain exon 40, it cannot accurately recapitulate the splice defect observed in patient cells. In contrast, a midigene construct that incorporated a larger *ABCA4* genomic region with exons 38 and 41 (Figure 2A) showed a similar aberrant splice pattern as the one described previously in patient cells (Aukrust et al. 2017) (Figure 2B). Further screening consisted of shorter AONs containing the sequence shared between AON31 and AON32 that were tested on midigene-transfected cells by transfection. Interestingly, this screening identified AON44, AON59 and AON60 as the most potent candidates that induced the greatest effect when compared to the initial molecule AON32 (Figure 2C). In addition, we noticed an increased rescue percentage when using the midigene model instead of the minigene; in fact, AON32 restored 46±3% of full-length *ABCA4* transcript in midigene-transfected cells, as opposed to the 20±0,5% of rescue detected in the minigene-transfected cells. This is probably driven by the high amount of double exon skipped *ABCA4* isoform that is expressed with the midigene but not with the minigene construct.

**Figure 2.**
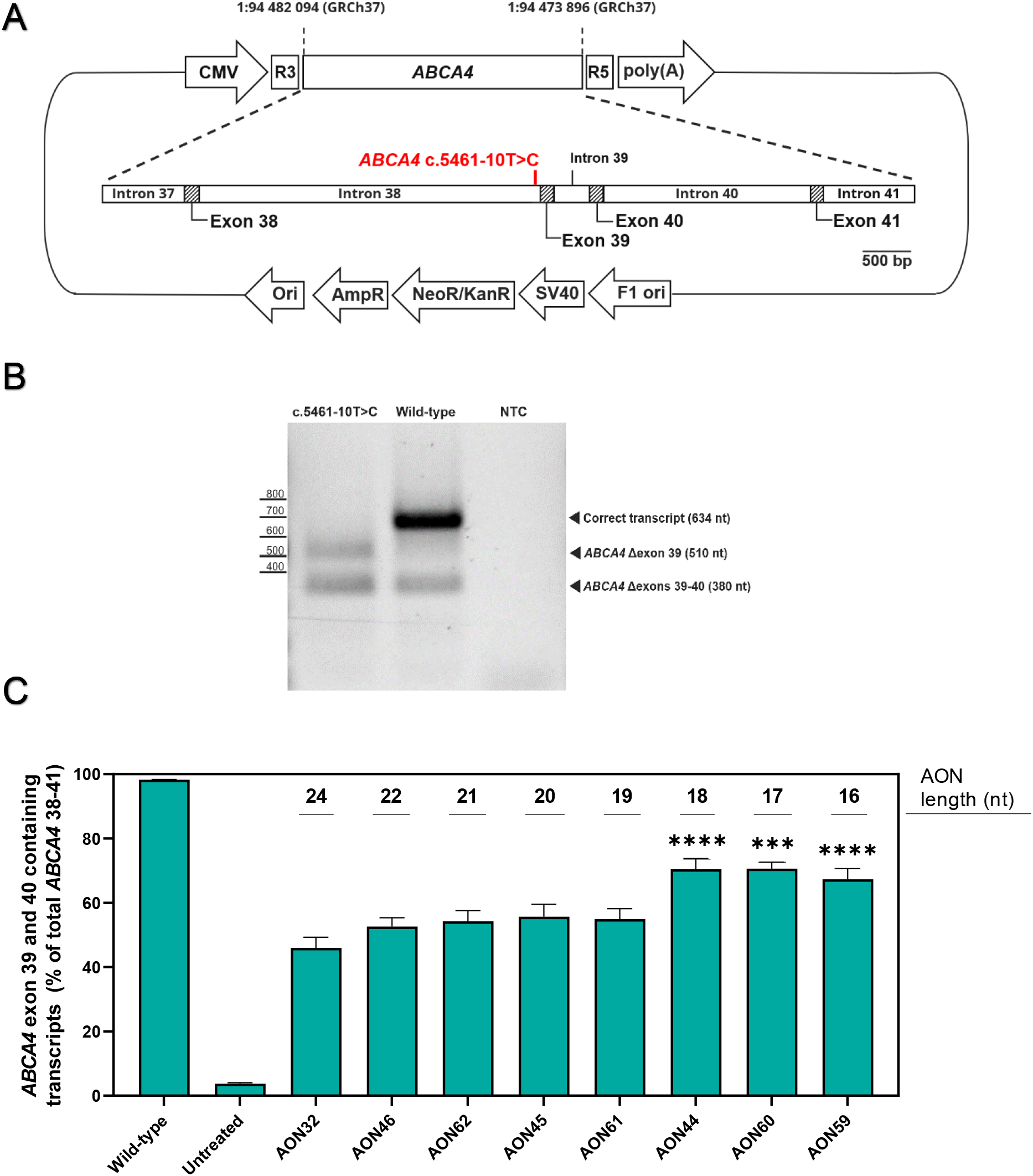
AON-induced *ABCA4* exon inclusion in splice-predictive midigene. (A) The midigene incorporates the *ABCA4* genomic region between intron 37 and 41, together with the *ABCA4* c.5461-10T>C mutation. The construct is flanked by Rhodopsin 3 and Rhodopsin 5 exons that contain strong splicing donor and acceptor sites, while the expression is initiated by the CMV promotor. To facilitate selection, the plasmids contain neomycin, kanamycin and ampicillin resistance genes. The midigene offers a few advantages over the minigene (Figure 7A): 1. because of a limited genetic environment, the double exon skip can’t be observed in the minigene; 2. since it carries a wider *ABCA4* sequence, the midigene produces *ABCA4* isoforms at more comparable ratios found in the native human retina with the *ABCA4* c.5461-10T>C mutation (B and Figure 3C). (B) The expression of the *ABCA4* c.5461-10T>C midigene in HEK293 cells transcribes in two truncated *ABCA4* isoforms: the *ABCA4* Δexon39 and *ABCA4* Δexons 39-40. The full length *ABCA4* was not detected, while the wild-type midigene displayed mostly expression of the correct transcript with traces of the double skip isoform. (C) The last AON screening with lead candidate AONs applied on midigene transfected cells demonstrated a clear effect over the untreated sample. HEK293 cells were treated with oligos using a 100 nM dose and the intake was facilitated with a transfection reagent. Twenty-four hours post-treatment, all applied oligos showed a significant effect over the untreated mutant sample. The AON32 is the longest AON construct that underperformed when compared to its shorter versions AON44, AON59 and AON60 that reached transcript correction at 71±3%, 67±3% and 71±2%, respectively. In addition, these 3 AONs showed significantly higher effect over AON32, unlike other oligos used in the experiment. These results determined that shorter AONs are likely more effective because of their easier cell intake in comparison to their longer versions. Data are shown as mean ± s.e.m., n=6, ***p<0.001, ****p<0.0001, ordinary one-way ANOVA test followed by Tukey’s multiple comparison test.

The dose-dependent effect of AON44, AON59 and AON60 was assessed following gymnotic administration (Figure S2). Here, the effect on splicing restoration was proportional to the concentrations of all AONs in treated samples. These experiments confirmed that shorter AONs (AON44, AON60 and AON59) could induce more splicing recovery when compared to the longer AON versions (AON32).

In addition, these 4 AON candidates were screened for pro-inflammatory potential in vitro, using a human peripheral blood mononuclear cell (PBMC) activation assay (Figure S3). The exposure of AONs at concentrations of 1 and 10 μM revealed a slight dose-dependent influence on cytokine release, which was comparable between candidates. Increases in cytokine secretion only reached statistical significance for MIP-1β, following exposure to AON44 [10 µM], and IL-6, following exposure to AON59 [10 µM]. We observed no effect on viability of PBMCs with any of applied treatments (data not shown).

### Application of AON treatment on CRISPR-Cas9 edited STGD1 ROs

Before assessing the AON treatment efficiency on ROs, we investigated the morphology and RNA content in CRISPR-Cas9 edited organoids homozygous for the c.5461-10T>C variant compared to the parent isogenic wild-type ROs. STGD1 and wild-type ROs were differentiated following the protocol published by Hallam et al. (2018) and their morphology and transcript content were examined 120 ± 3 days post-differentiation. Both groups showed a phase-light neural retina region at their edges, which is characteristic light microscopic appearance for ROs older than 90 days (Figure 3A). In addition, we observed a short “brush border” surrounding the margins of both groups; this contains presumptive inner and outer segments of photoreceptor cells that is expected to emerge after 120 days after differentiation. Transcript analyses revealed similar expression levels of the photoreceptor precursor gene *CRX*, retinal identity marker genes (*NRL* and *NR2E3)*, and *USH2A* (Figure 3B). Interestingly, the expression of *ABCA4* was significantly reduced in STGD1 organoids (Figure 3B and Table S2), indicating the possible presence of nonsense-mediated decay that is activated by the out-of-frame RNA transcripts in STGD1 organoids. In wild-type retinal organoids, *ABCA4* was not detected before day 35 post-differentiation; the expression steeply increased until day 120 (Figure S4B), after which the increases were more gradual.

**Figure 3.**
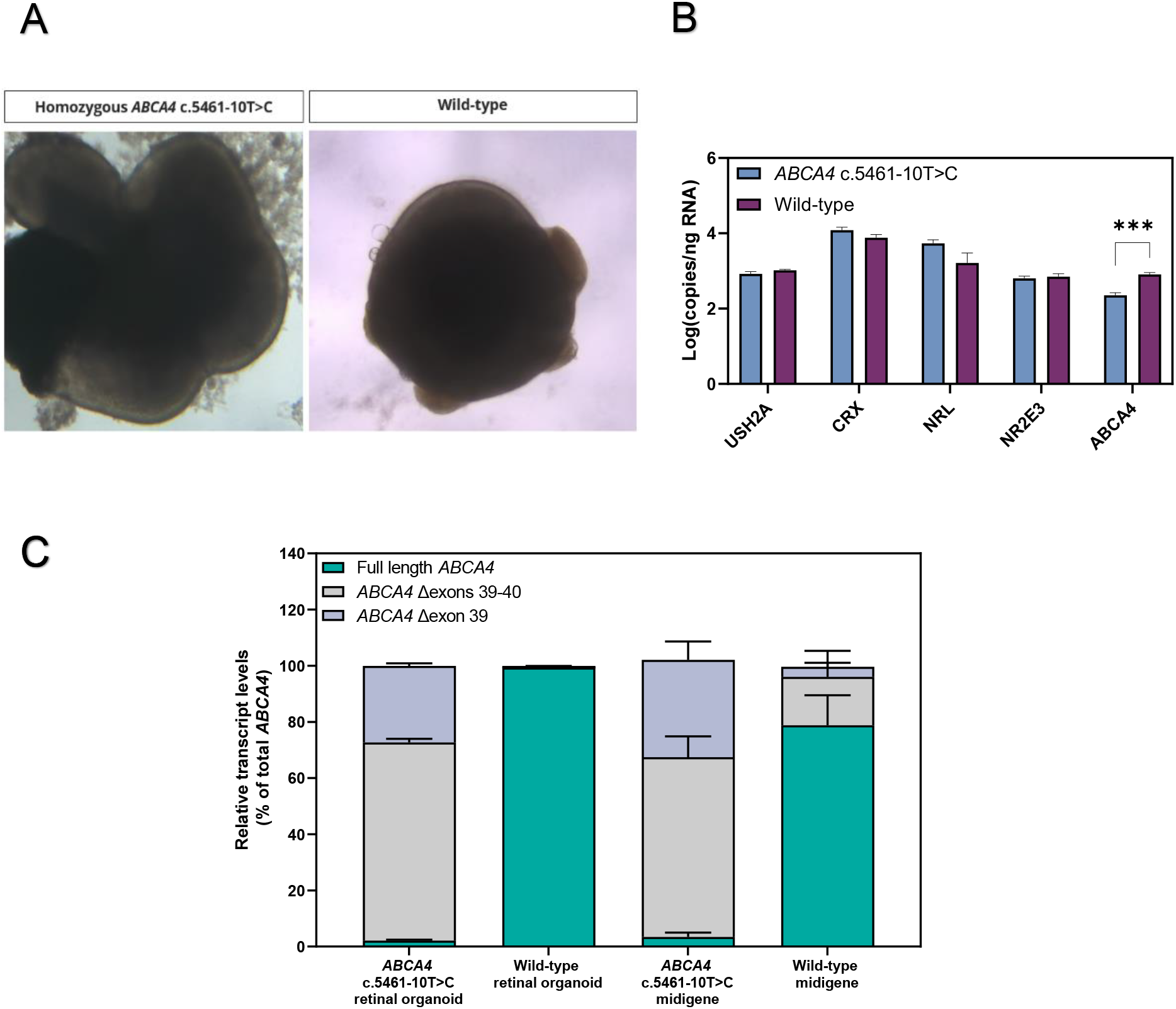
Morphological and transcript comparison of wild-type and gene edited homozygous *ABCA4* c.5461-10T>C retinal organoids. (A) Morphology assessed at day 120 of organoid differentiation suggests ROs derived from homozygous c.5461-10T>C and the control isogenic parent cell line displayed neural retina (thin light rim at the margin of the ROs) characteristic for this stage in organoid development. Moreover, both groups showed the development of the brush border with photoreceptor cells above the neural retina, which is expected to develop after day 120 of the organoid differentiation. (B) These ROs were analyzed for the total transcript expression of photoreceptor precursor gene (*CRX*), photoreceptor markers (*NRL, NR2E3* and *USH2A*) and *ABCA4*. *CRX* and the photoreceptor markers were expressed similarly in both groups. *ABCA4* was expressed at higher levels in wild-type organoids, suggesting a possible presence of nonsense-mediated decay and a consequent degradation of out-of-frame transcripts in *ABCA4* c.5461-10T>C organoids. A multiple t-test was conducted to confirm the similarities in RNA content in wild-type and STGD1 ROs reported in Table S5. Data are shown as mean ± s.e.m., n=4. (C) Transcript isoform comparisons show that c.5461-10T>C ROs and midigene have similar content of full-length *ABCA4* isoform and isoforms missing exon 39 or exons 39 and 40. On the other hand, the wild-type organoid displayed almost exclusively full-length *ABCA4* mRNA, whereas the wild-type midigene has higher levels of both *ABCA4* Δexon39 and *ABCA4* Δexon39-40. Data are shown as mean ± s.e.m., n=3. ***p<0.0001, ordinary one-way ANOVA test followed by Dunnet’s multiple comparison test.

We compared the presence of the different *ABCA4* splice forms between ROs and midigenes. The c.5461-10T>C variant led to similar ratios of *ABCA4* transcripts in both organoid and midigene models: exon 38-39-40-41 isoforms were barely detected, while the Δexons39-40 isoform was the most prominent. In contrast, the wild-type ROs contained almost exclusively the correct transcript, unlike the wild-type midigene that displayed some missplicing (Figure 3C).

CRISPR-Cas9 edited ROs homozygous for c.5461-10T>C were treated gymnotically with AON32, AON44 and AON60 once they reached 150 days of age. The treatment followed a ‘wash-out’ regimen for 4 weeks: here, the AONs were added only on day 0 of treatment at a 1.5 µM concentration, and at each 50:50 media change the AON concentration would be halved. This study included the AONs with the 2’OMe modified sugar rings tested previously in cells; in addition, we included the same AON sequences carrying the 2’MOE sugar modifications to examine the possible difference in therapeutic effect between the two chemistries. Isoform-specific analysis revealed that AON60 and its 1-nt longer version AON44 reached 35±4% and 33±7% of transcript correction (Figure 4A). The longest molecule AON32 showed no significant improvements compared to the scrambled sample, and the two different sugar chemistries showed only small differences in outcome. Because of its superior theoretical parameters, AON44 2’MOE was selected for further analysis in ROs and named QR-1011.

**Figure 4.**
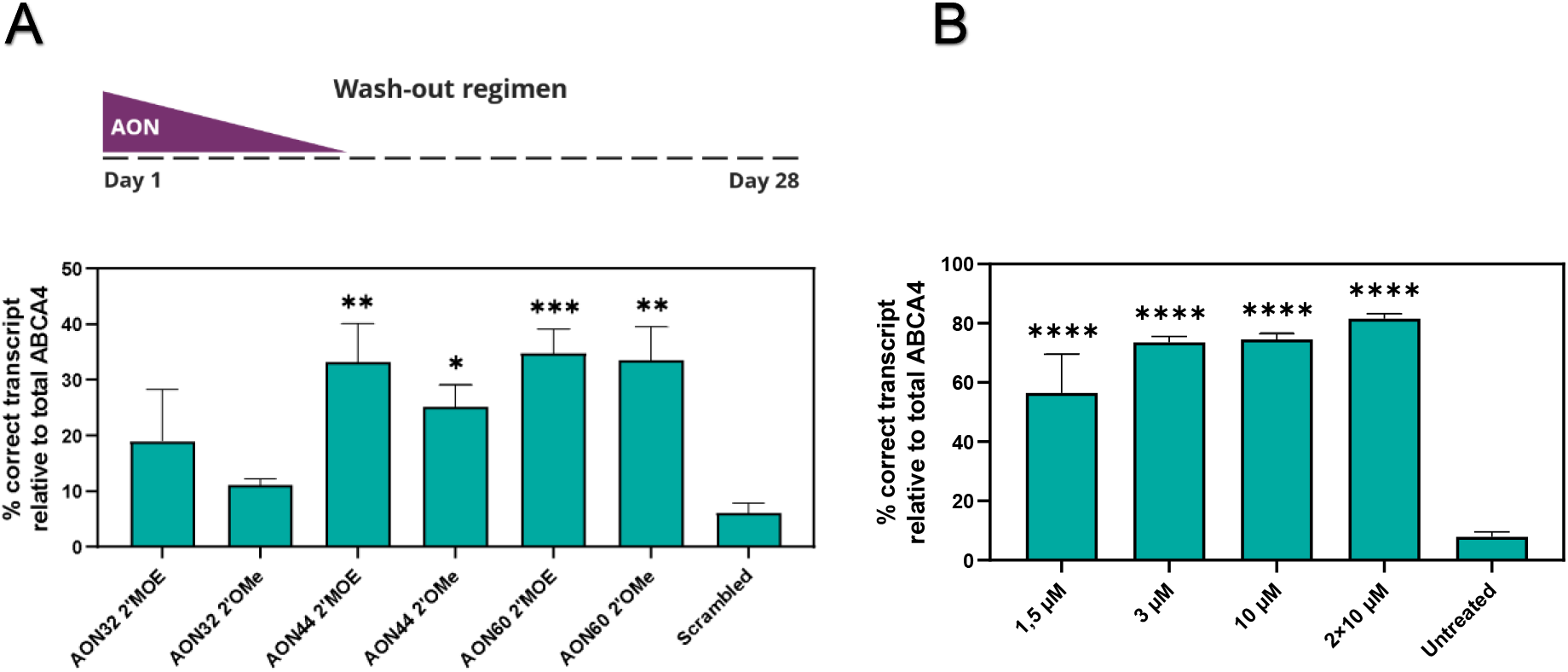
AON treatment of gene edited and patient-derived homozygous c.5461-10T>C ROs show high levels of rescued *ABCA4* transcript. (A) AON gymnotic treatment of 150 day old gene edited ROs at a 1.5 µM dose of lead candidates AON44 and AON60, together with the longer version AON32. In addition to the 2’OMe chemistry used previously in cells, the 2’MOE chemistry was included in all three AONs to evaluate differences in therapeutic effect of the two chemical modifications. Upper panel shows the 4-week long treatment used a wash-out regimen in which the oligo was added only at day 0 of treatment and its concentration was halved by each medium change. Even though AON60 showed slightly higher modulation effect when compared to AON44, this last was selected as best lead candidate due to the more stable parameters of the molecule and was consequently named QR-1011. (B) QR-1011 was administered to patient-derived homozygous c.5461-10T>C ROs when they reached 180 days of differentiation. The therapeutic effect of the molecule was investigated at 4 different concentrations: 1.5, 3 and 10 µM, with all concentrations added once at day 0, and only the highest dose was again administered at treatment day 14. The wild-type isoform in the untreated samples was present at <8%. Data are shown as mean ± s.e.m., n=6 per condition. Asterisks display the significant differences with the control group treated with scrambled oligo (*p≤0,05, **p≤0,01, ***p<0,001, ****p<0,0001, ordinary one-way ANOVA test followed by Dunnet’s multiple comparison test).

### In silico analysis does not show any relevant off-target effect of QR-1011

A search against the RefSeq database showed that QR-1011 has no full complementarity to any targets in human mRNA and DNA, apart from the intended target in *ABCA4*. We also identified no target with one mismatch, whereas one coding region in *MGRN1* showed to be a potential off-target with two mismatches (GRCh37, chr16:4683901-4683916). In addition, potential off-targets in genomic DNA with 2 mismatches were predicted for 5 intergenic and 16 deep-intronic regions (Table S4); since the distance of the flanking exons for all genes was ≥ 225 bases, it was concluded that the possible interference with splicing of these genes by QR-1011 was unlikely (Liu and Zack 2013). Given the short size of QR-1011, a near perfect match would be necessary for efficient hybridization and it is therefore highly unlikely that the oligo would efficiently hybridize to targets with ≥ 2 mismatches, as reported previously (Garanto et al. 2019).

### Range of activity of lead candidate QR-1011 in patient-derived ROs homozygous for *ABCA4* c.5461-10T>C

Based on previous unpublished studies, we noticed that longer treatment periods up to three months enable more AON-mediated transcript correction, potentially related to endosomal storage and release of AON, that could consequently lead to the production of more wild-type protein. Hence, the next study consisted of a wash-out treatment 8 weeks long where QR-1011 was applied at 4 different concentrations. Homozygous c.5461-10T>C patient-derived iPSCs (Sangermano et al. 2016) were differentiated to ROs, QR-1011 was administered at D180 at concentrations of 1.5 µM, 3 µM, and 10 µM and the organoids were harvested 56 days later. To explore the relative efficacy of QR-1011 dosing and wash out, some of organoids treated with the 10 µM dose were retreated with another 10 µM dose 14 days after the first dose and the treatment continued for 6 additional weeks. The isoform analysis revealed that the 1.5 µM dose of QR1011 reached 56±13% correct exon 38-39-40-41 *ABCA4* transcript of the total detected *ABCA4* (Figure 4B). Interestingly, the organoids that underwent the 10 µM and the 2×10 µM treatments did not show significantly higher restoration of splicing when compared to the group treated with 3 µM QR-1011. Indeed, post-treatment analysis of RNA indicated that the AON activity at 3 µM restored 74±2% of correct splicing and reached a plateau phase, suggesting this dose potentially corrected most of the available aberrant transcript. This experiment also showed that an 8-week long treatment induced more AON-mediated correction when compared to a 4-week long treatment; moreover, the 1.5 µM dose here induced 41% more correctly spliced *ABCA4* than the previous experiment where the same concentration of AON was administered (Figure 4B). In addition, we noticed that the untreated patient-derived ROs displayed slightly higher, yet not significantly increased amounts of correct transcript (8±2%), compared to those detected previously in the CRISPR-Cas9 modified organoids (6±2%).

### Patient-derived ROs display a dose-response splicing rescue of RNA that correlates with wild-type ABCA4 protein rescue

To assess the activity of QR-1011 at lower concentrations, patient-derived ROs were treated with QR-1011 at concentrations of 0.375 µM, 0.75 µM, 1.5 µM and 3 µM. The treatment followed the same design applied in the previous experiment described above. This treatment included two positive control groups of wild-type ROs that were either treated with a scrambled AON or left untreated. The aim of this experiment was to compare the restoration of *ABCA4* levels to those observed in wild-type ROs, in contrast to in previous treatments where the AON effect was calculated as part of total *ABCA4* within the same sample. Eight weeks post-treatment the *ABCA4* 38-39-40-41 transcript content showed a clear dose-response in patient-derived ROs; an increase of 21±1.5% in the generation of correct *ABCA4* isoform was detected with the lowest 0.375 µM dose of QR-1011, whereas the highest 3 µM dosage rescued 53±5% of the correct transcript when compared to the total *ABCA4* content detected in untreated wild-type ROs (Figure 5A).

**Figure 5.**
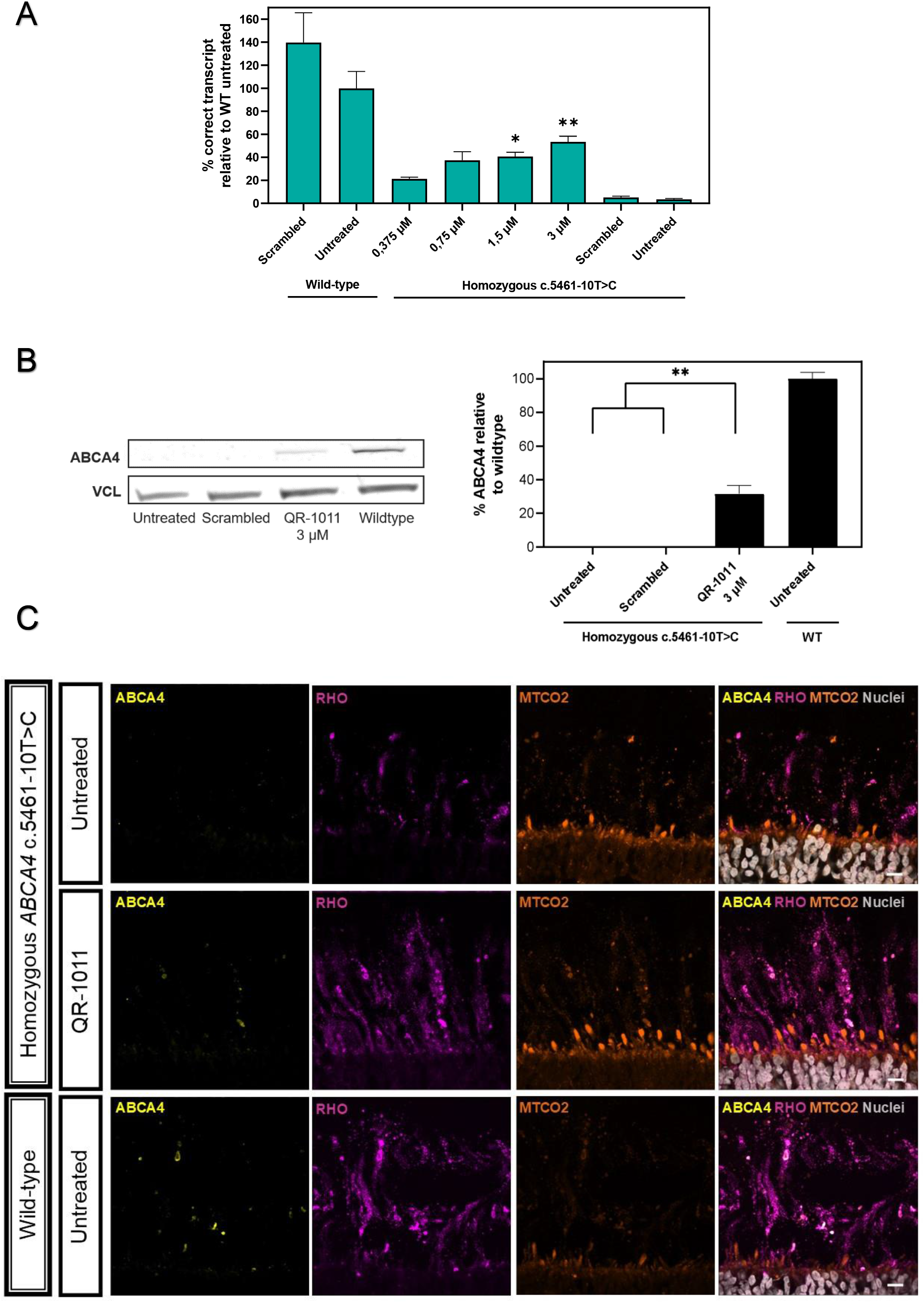
Low concentrations of QR-1011 have high restoring activity on RNA splicing and rescue of the wild-type protein in patient-derived c.5461-10T>C ROs. (A) Splice adjustment efficacy with clinically relevant dosages of QR-1011 in ROs deriving from a biallelic c.5461-10T>C patient cell line. Four different concentrations of QR-1011 were administered to patient-derived ROs (n=6); in addition, a 3 µM dose of scrambled AON was given to both c.5461-10T>C and wild-type organoids as a negative control. After a 56-day treatment, all patient-derived samples showed the splice restoring activity of QR-1011; in addition, the scrambled AON showed no effect on ABCA4 splicing in homozygous ROs. These contained almost only misspliced *ABCA4* isoform, as opposed to the wild-type ROs that served as positive controls. Data are shown as mean ± s.e.m., n=6 for all conditions. (B) Western blot analysis (n=3) identified significant levels of rescued ABCA4 protein in treated ROs. The expression of ABCA4 was determined with the anti-ABCA4 clone 5B4 antibody, while vinculin (VCL) was used as a loading control. Untreated c.5461-10T>C ROs and those subjected to treatment with scrambled AON contained no detectable protein. All samples were normalized to the average signal obtained from wild-type organoids (control). Data are shown as mean ± s.e.m., n=3. *p<0,05, **p<0,01, ordinary one-way ANOVA test followed by Dunnet’s multiple comparison test. (C) ABCA4 protein immunoreactivity (yellow) in treated patient-derived organoids colocalized within the outer segments (OS) stained with anti-rhodopsin 4D2 clone (magenta) of photoreceptor cells and resembled the localization found in wild-type organoids. The inner segments (IS) were visualized with the mitochondrial-targeting antibody MTCO2 (orange). ABCA4 was visualized with anti-ABCA4 clone 3F4 targeting the C-terminal end (yellow) and DAPI nuclear staining is shown in grey.

To assess if the presence of an off-target oligo interferes with the transcript content, we investigated whether the wild-type untreated ROs and those that underwent treatment with the scrambled AON showed significant difference in their total expression of *ABCA4*. Predictably, statistical analysis of these two groups did not indicate any significant difference in the expression or splicing of *ABCA4*.

Western blot analysis was performed on a pool of patient-derived ROs treated with 3 µM QR-1011 and compared to ROs that were untreated or treated with scrambled AON (used as negative controls) and untreated wild-type ROs that served as a positive control. The antibody directed against the N-terminus of the protein revealed AON-induced rescue of the wild-type protein in bi-allelic variant ROs; these expressed 32±5% of newly generated protein relative to the wild-type organoids (Figure 5B). ABCA4 protein was undetectable in untreated homozygous c.5461-10T>C ROs, correlating with the very low levels of the correct in-frame transcript.

To investigate the trafficking and the subcellular expression of the newly generated ABCA4 protein in more detail following AON treatment, cryosections of ROs were investigated by immunohistochemistry. The fragile outer segments of photoreceptor cells surrounding the ROs were preserved with gelatin-embedding, as described previously (Cowan et al. 2020). Wild-type ABCA4 was visualized by immunofluorescence with an ABCA4 antibody targeting the C-terminal part of the protein. We observed that in wild-type ROs ABCA4 immunoreactivity was exclusively in the outer segments of the photoreceptor cells co-stained with rhodopsin antibody. Interestingly, 3 µM treated patient-derived ROs displayed ABCA4-immunoreactivity in the photoreceptor outer segments, as opposed to the untreated patient ROs where no immunoreactivity was detected (Figure 5C). This staining confirmed that the trafficking of the rescued ABCA4 protein is in line with what is observed in the native human retina. We did not detect any protein retention in the inner segment of photoreceptor cells that was stained with a mitochondria-targeted antibody against MTCO2. The same localization of ABCA4 immunoreactivity was observed in wild-type ROs. These findings suggest that the AON-mediated treatment not only restores wild-type ABCA4 protein, but the protein generated upon AON treatment is also trafficked to the expected subcellular compartment.

## DISCUSSION

In this study, we report the development and validation of target-specific antisense oligonucleotides as a possible therapeutic approach to correct the aberrant splicing of *ABCA4* caused by the severe STGD1-causing variant c.5461-10T>C. This variant was previously identified as a non-canonical splice site variant that induces generation of deleterious transcripts (lacking either the single exon 39 or exons 39 and 40) that lead to frame-shifts in the open reading frame. Extended *in vitro* AON screenings in a c.5461-10T>C midigene model identified potent lead candidates that were able to correct 70% of the aberrant splicing. These molecules were validated in differentiated ROs where we observed high levels of splicing correction accompanied by rescued wild-type ABCA4 protein.

AONs have demonstrated promising results in several pre-clinical studies for IRDs, by correcting pathological RNA processing events associated with entire exon skipping, complete degradation of abnormal transcripts or pseudo-exon exclusion. In the clinic, AON-mediated splicing therapy was well tolerated and able to significantly improve the best corrected visual acuity (BCVA) in phase 1 clinical trials with Sepofarsen (LCA10) (Cideciyan et al. 2021; Russell et al. 2022) and Ultevursen (USH2A associated RP and Usher syndrome) (Dulla et al. 2021). These AONs delay disease progression through pseudo-exon and in-frame exon skipping, respectively. The efficacy of both compounds is currently being investigated in phase 2/3 clinical trials.

Considering that ABCA4 protein is a complex membrane protein constituted of 12 transmembrane helixes that are involved in the transport of substrates (Xie et al. 2021), many of the disease-causing variants are predicted or known to lead to protein misfolding (Molday et al. 2021). Splicing variants in *ABCA4* are estimated to comprise 25% of all STGD1-causing variants, which emphasizes the importance of the advancement of AON-mediated therapy because of its splicing-manipulating activity. Here, AON-mediated exon inclusion was implemented, for the first time, to our knowledge, in IRD treatment development, to re-include skipped exons 39 and 40 in *ABCA4* and correct the disrupted splicing caused by *ABCA4* c.5461-10T>C. This AON mechanism of action could be of major importance in the development of therapies for STGD1, since the complexity of the ABCA4 protein structure suggests that it is unlikely that major truncation of the original amino acid sequence would be tolerated without leading to a disease phenotype (Bauwens et al. 2019). The potential of AON-based re-inclusion of skipped exons as therapy has been illustrated by several pre-clinical studies for Pompe disease (van der Wal et al. 2017), cystic fibrosis (Igreja et al. 2016) and Alzheimer’s disease (Hinrich et al. 2016). All reported a significant post-treatment functional rescue that is a prerequisite for ameliorating disease-associated phenotypes. The most effective AONs from these studies block intronic splice silencers (ISSs) located in the adjacent introns to promote the exon inclusion. Similarly, QR-1011 is designed to block three strong ISSs located in intron 39 of *ABCA4* and restore the canonical splicing (Figure S1C).

*ABCA4* c.5461-10T>C is considered the most common severe variant that underlies STGD1 (Cornelis et al. 2022). Even though the clinical features associated with STGD1 reveal a wide range of heterogeneity, the severity of *ABCA4* variants is directly correlated with the onset of the disease (Fakin et al. 2016).The -10T>C variant is more often found in combination with one other moderate or mild variant in ABCA4 than in a homozygous state (Cornelis et al. 2022; Maugeri et al. 1999). Considering that severe mutations lead to the early STGD1 onset where the progress of the disease is faster, and milder mutations are associated with slower development of STGD1 hallmarks (Fujinami et al. 2015) (Cremers et al. 2020), the advanced stage of STGD1 could be significantly postponed or even completely repressed by correcting the aberrant splicing due to c.5461-10T>C. In the case of two deleterious *ABCA4* alleles, the disease-associated changes characteristic of the early stage are observed in the region limited to the macula. The most progressed phases of the disease exhibit severe degenerative lesions of the retina that extend across the posterior pole of the retinal tissue and cause severe visual impairment. The loss of photoreceptor cells in STGD1 prevents the reversion of the disease-associated phenotype; however, individuals diagnosed in an early stage of the disease could benefit greatly from the AON-based intervention since this would slow or stop the progress of advanced STGD1 features. Proof of concept acquired from patient-derived ROs show that QR-1011 was effective in restoring correct *ABCA4* transcript splicing followed by production of wild-type protein. Regarding its safety profile, high concentrations of QR-1011 did not exhibit overt toxicity throughout the screenings in cells and studies in ROs. The modest influence on secretion of pro-inflammatory cytokines and chemokines upon exposure to PBMCs suggests a favorable immunostimulatory profile. Nevertheless, further dedicated toxicology studies to completely assess potential adverse effects of the molecule will be required. On the other hand, low QR-1011 concentrations, ranging from 0.375 and 3 µM, were able to correct the transcript and higher amounts of aberrant splicing correction were observed in longer organoid treatments, compared to shorter treatments. We noticed that 4-week long treatments were less effective than 8-week long treatments even with the same AON concentrations. This suggests that the therapeutic effect does not stop soon after exposure to QR-1011, but is rather more durable and might prolong the interval between the doses once in clinic. The likely mechanism behind this event involves the slow endosomal release of the AON to the nucleus after its entrapment in early or late endosomes upon endocytosis (Juliano et al. 2014). Future studies on non-human primates will be required to further assess the tolerability and the safety profile of QR-1011, which can determine the effective clinical dose for intravitreal delivery. These should be accompanied by investigations of the pharmacokinetic properties of the molecule in order to establish the optimal dosing interval.

The characterization of disease-associated clinical features and investigation in novel therapies demand robust and credible pre-clinical models. ROs differentiated from pluripotent stem cells have demonstrated promising *in vitro* applications; several groups reported protocols for generation of 3D layered retina-like tissues (Afanasyeva et al. 2021). To evaluate the splice-regulating effect of our AONs, we used CRISPR-Cas9 edited and patient-derived ROs bi-allelic for *ABCA4* c.5461-10T>C. These STGD1 organoids were differentiated using two different protocols published by Hallam et al. (2018) and Hau et al. (2022). Once they reached the mature stage 3 after 120 days of differentiation, the ROs appeared similar, despite differences in differentiation procedures. They had similarities in lamination accompanied by a developed surrounding brush border. In addition, both protocols produced ROs that morphologically resembled the wild-type ROs, despite the presence of the severe *ABCA4* variant on both alleles. The RNA content of most retinal markers was concordant between all types of ROs, with exception of *RHO* that was detected at significantly higher levels in patient-derived and controls ROs produced with the adherent non-embryoid body method (Hau et al) than in gene-edited and control ROs produced with the Hallam et al method. In addition, we observed a reduced level of *ABCA4* transcript in STGD1 organoids, as opposed to wild-type ROs, which is most likely due to nonsense-mediated decay of the exon-skipping transcripts in STGD1 organoids. To study the expression and localization of the wild-type protein in detail, we conducted protein analysis by Western Blot and Immunohistochemistry. Considering the fragile nature of the photoreceptor outer segments we used embedding in gelatin, which previous studies reported improved the preservation of structures surrounding the ROs (Cowan et al. 2020). ABCA4 protein was clearly expressed in wild-type ROs and its localization was confined to the outer segments of retinal photoreceptors, as observed in the human retina. ABCA4 was much less present in the outer segment when compared to rhodopsin, which is in line with earlier observations conducted in mouse rod cells where the molar ratio of ABCA4 to rhodopsin was 1:300 (Skiba et al. 2021). The protein assays in STGD1 ROs that underwent the AON treatment confirmed the utility of ROs as *in vitro* model for STGD1; we confirmed that the rescued protein is trafficked and co-localizes in outer segments of photoreceptor cells. As reported previously, the correct subcellular localization of rescued protein is important for alleviation of the STGD1 phenotype (Liu et al. 2019). The amount of ABCA4 residual activity required to prevent the STGD1 phenotype in affected individuals remains unclear, although previous inquiries on *abca4*^-^/^-^ mice identified significantly reduced traces of deposited lipofuscin after compensating just 10% of wild-type protein (Tornabene 2019). In addition, the activity of isolated and purified protein derived from wild-type and several disease-causing missense variants has been determined (Pollock et al. 2014; Quazi and Molday 2013); these studies suggest the basal activity of severe variants to be < 25 % of the wild-type protein (Molday et al. 2021). Importantly, we quantified the restored ABCA4 protein post-treatment with QR-1011 at 32±5% of the wild-type levels, as opposed to untreated bi-allelic c.5461-10T>C samples where no protein was detected.

In conclusion, QR-1011 showed robust therapeutic potential by correcting high levels of truncated transcripts in *ABCA4* c.5461-10T>C when administered to both midigene-transfected cells and 3D human ROs. In 8-week long treatment periods, the AON-corrected transcripts led to the production of wild-type ABCA4 protein, which trafficked to the outer segments of photoreceptor cells in ROs. The measured amounts of rescued RNA and protein suggest the AON effect would be sufficient to alleviate the STGD1 phenotype, and therefore QR-1011 shows potential as a therapeutic strategy for the most common severe STGD1-causing variant *ABCA4* c.5461-10T>C.

## MATERIALS AND METHODS

### Generation of *ABCA4* wild-type and *ABCA4* c.5461-10T>C minigenes and midigenes

pIC-neo.Rho3-5.MCS (Gamundi et al. 2008) was generated by replacing the USH2A sequences of pCI-neo.Rho.USH2A-PE40-wt by a custom MCS containing the following restriction enzyme recognition sites: XhoI, EcoRI, MluI, EcoRV, XbaI, SalI and Cfr9I designed to aid in downstream cloning steps. The custom MCS was generated by annealing DNA oligonucleotides 5’-CTCGAGAATTCACGCGTGGTGATATCACCTCTAGAGTCGAC-3’ and 5’-CCCGGGTCGACTCTAGAGGTGATATCACCACGCGTGAATTCT-3’. The resulting fragment was used in a ligation mixture together with the backbone plasmid, digested with XhoI and Cfr9I (Thermo Fisher Scientific).

To generate a *ABCA4* c.5461-10T>C minigene, the pCI vector backbone and a synthetic dsDNA sequence (gBlock; Integrated DNA Technologies) containing the *ABCA4* minigene genomic region and the c.5461-10T>C mutation were digested using the *Eco*RI (New England Biolabs) and *Sal*I (Thermo Fisher Scientific). The digested vector was loaded on 1% agarose gel, isolated and purified using the Nucleospin Gel and PCR Clean-up kit according to the manufacturer’s instructions (Bioké). The digested gBlock was purified directly using the same kit. Digested fragments were ligated overnight at 16°C with T4 ligase (Thermo Fisher Scientific) following the manufacturer’s protocol. The ligation reaction was used to transform DH5α competent cells (Thermo Fisher Scientific) according to manufacturer’s protocol.

To generate a *ABCA4* wild-type midigene, genomic DNA from HeLa cells (ATCC) was extracted with the DNeasy Blood & Tissue Kit (QIAGEN). The *ABCA4* genomic region between intron 37 and intron 41 was amplified with primers (Integrated DNA Technologies) containing recognition sites for *Eco*RI and *Sal*I, using the Phusion™ High-Fidelity DNA Polymerase kit (Thermo Fisher Scientific) according to the manufacturer’s protocol. The wild-type PCR product and the digested pCI vector backbone were ligated and transformed into GT115 competent cells (InvivoGen). To introduce the *ABCA4* c.5461-10T>C mutation, the wild-type plasmid and a gBlock (Integrated DNA Technologies) from intron 37 to intron 41 containing the c.5461-10T>C mutation were digested with *Box*I and *Bsi*WI (Thermo Fisher Scientific) and, subsequently, ligated and transformed into GT115 competent cells.

### AON screening in HEK293 cells

HEK293 cells (ATCC) were cultured with Dulbecco’s Modified Eagle Medium (DMEM; Gibco) with 10% Fetal Bovine Serum (Biowest, France) at 37°C with 5% CO_2_. 2 × 10^5^ cells were transfected with 50 ng of either *ABCA4* minigenes or midigenes with Lipofectamine 3000 Transfection Reagent (ThermoFisher Scientific) by following the manufacturer’s protocol. The AON was delivered by transfection or gymnotically as described by Dulla et al (2021). The RNA was extracted 48h post-treatment with RNeasy Plus Mini Kit (QIAGEN, Germany) and 500 ng was reverse transcribed using the Verso cDNA Synthesis Kit (ThermoFisher Scientific) by following the manufacturer’s protocol. 5 ng of cDNA was analyzed with isoform-specific droplet digital PCR assays and ddPCR Supermix for probes (Bio-Rad). Primers and probes (Integrated DNA Technologies) used for dPCR are listed in Table 2. The following PCR program was used: enzyme activation at 95°C for 10 minutes (1 cycle), denaturation at 95°C for 30 seconds and annealing/extension at 60°C for 1 minute (40 cycles), and enzyme deactivation at 98°C for 10 minutes (1 cycle). The fluorescence signal of individual droplets was measured in the QX200™ Droplet Reader (Bio-Rad). In each experiment the thresholds to separate the positive droplet population from the negative were set manually. The following formulas were used to calculate the percentage of *ABCA4* exon 39-40 inclusion isoform in minigene:

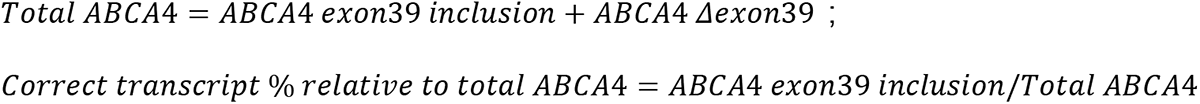

The formulas that were used in the midigene model and ROs were the following:

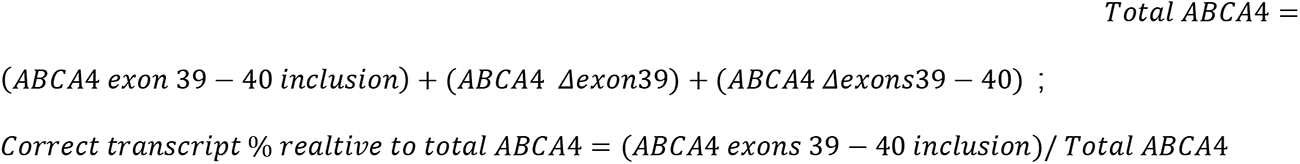

The mean percentage of detected full-length *ABCA4* isoform post-treatment was statistically analyzed using GraphPad Prism 9 with ordinary one-way ANOVA test followed by Tukey’s multiple comparison test.

### In-silico analysis of possible off-target effect of selected lead AON molecules

*Homo sapiens* RefSeq RNA was downloaded from the NCBI website and used for finding potential off-targets of QR-1011 (https://ftp.ncbi.nlm.nih.gov/refseq/H_sapiens/mRNA_Prot/). *Homo sapiens* gene sequence database was generated from human genome sequence database (https://ftp.ensembl.org/pub/release-90/fasta/homo_sapiens/dna/) using Ensemble gene annotations (https://ftp.ensembl.org/pub/release-90/gff3/homo_sapiens/). *Homo sapiens*mRNA and pre-mRNA sequence databases were queried using a custom made bioperl program. The reverse complement sequence of QR-1011 (CCGAGGCCCATGGAGCAT) was used for the searches. In cases where QR-1011 matched to multiple mRNA isoforms of the same gene, it is considered as one match/target and match with highest complementarity (least mismatches) is reported. Ensemble Genome Browser (http://www.ensembl.org/index.html) was used to find the exact location of the match and to calculate the distances to the flanking exons. Target gene was queried against human sequences. From the results, the most abundant transcript was selected and target sequence was searched.

### Assessment of the immunostimulatory and cytotoxic potential of lead AON candidates AONs in PBMCs

Buffy coats, the fraction of an anti-coagulated blood sample that contains most of the white blood cells and platelets following centrifugation of the blood (500 mL blood in 70 mL citrate phosphate dextrose coagulant), from 5 healthy human (consensual) blood donors, were obtained from Sanquin Blood Supply in Rotterdam, the Netherlands. PBMCs were isolated from each buffy coat within 24 h after blood collection, aliquoted and cryopreserved. PBMCs were stimulated for 48 hours with QR-1011 candidates AON32, AON44, AON59 or AON60 at a concentration of 1 μM and 10 μM; positive control R848 (1 μM); or PBS (vehicle control) at 37°C under a 5% CO2 atmosphere. For every donor, all conditions were tested in triplicate in 96-well round-bottom microtiter plates. The total number of viable PBMCs per well was 3·10^5^. R848 (Resiquimod; InvivoGen; tlrl-r848), a potent Toll-like receptor (TLR)7/8 agonist, was selected as a positive control for its strong and robust immune-activating properties, inducing the production of pro-inflammatory cytokines. Also, R848 acts upon the TLRs that are most likely to be involved in recognition of single-strand RNA, arguably making it the most relevant positive control for this purpose. After incubation, cell culture supernatant was isolated following centrifugation (300 relative centrifugal force [RCF], 5 min, room temperature). Viability of PBMC following exposure to test items was assessed by resazurin reduction assay (CellTiter-Blue Reagent, Promega, Madison, WI, USA). Cytotoxicity was assessed by measurement of lactate dehydrogenase in the cell culture supernatant (CyQUANT™ LDH Cytotoxicity Assay, ThermoFisher, Waltham, MA, USA). Readout of viability and cytotoxicity assays was performed on a SpectraMax M5 Microplate reader. Cytokine levels in PBMC culture supernatants were measured using the MILLIPLEX MAP Human Cytokine/Chemokine Magnetic Bead Panel-Custom 6 Plex-Immunology Multiplex Assay (Millipore; #HCYTOMAG-60K). Analytes included were the following: interferon (IFN)-α2, IL-6, IP-10, MIP-1α, MIP-1β, and tumor necrosis factor (TNF)-α. Assay plates were read on the Luminex MAGPIX platform (Luminex, San Francisco, CA, USA). Analysis of the Luminex data was performed in Bio-Plex Manager 6.1 software (Bio-Rad). Standard curves were fitted using 5 parameter logistic regression. Cytokine concentrations that were outside of the detectable range of the assay were imputed for the purpose of calculation and statistical analysis. Values below the limit of detection (LOD), rendered “out of range <” by the analysis software, were imputed with a concentration value of ½ ⋅ LOD. The LOD values, which were empirically determined by the manufacturer of the Luminex kit, were derived from the technical data sheet. Conversely, cytokine concentrations that were above the upper limit of quantification, rendered “out of range >” by the analysis software, were imputed with a concentration value of two times the concentration of the highest calibrator. Statistical analysis of the cytokine data was performed using GraphPad Prism 9 software. Prior to statistical comparison and graphical representation of cytokine secretion data, outlier removal was performed for every treatment condition except for positive control R848 using the “Identify Outlier” option, using ROUT method with a Q-value of 0.5%. Subsequently, log-transformed cytokine concentration values were subjected to matched comparison to PBS-treated controls using mixed-effects analysis, correcting for multiplicity using Dunnett’s correction.

### Generation of wild-type and homozygous *ABCA4* c.5461-10T>C ROs

GibcoTM Episomal hiPSCs line #A18945 was CRISPR-Cas9-edited by the manufacturer (ThermoFisher Scientific) to carry the *ABCA4* c.5461-10 T>C variant on both alleles. The wild-type and mutant iPSCs were cultured on Matrigel® hESC-Qualified Matrix coating (Corning, NY) with mTeSR1 medium (StemCell Technologies, Canada) and 1×mTeSR1 Supplement (StemCell Technologies). Patient derived homozygous c.5461-10T>C iPSCs were described previously (Sangermano et al 2016). The iPSCs were differentiated in ROs following the protocol described by Hallam et al (2018) or Hue et al (2022).

### AON treatment in ROs

ROs were treated with AONs after at least 150 days of differentiation. At treatment initiation, culture medium was fully removed and fresh medium containing AON was added. Every two days half of the culture medium was replaced by fresh culture medium, resulting in a gradual decrease in AON concentration in the medium. Eight weeks post-treatment the culture medium was removed, the ROs were washed in PBS and 300 µL of TRIreagent (Zymo Research, CA) was added. The samples were snap-frozen in liquid nitrogen and stored at -80°C until RNA extraction.

After thawing, the organoids were lysed by passing through a 25-gauge needle until homogenized (Henke Sass Wolfe), and the RNA was extracted with the Direct-Zol RNA MicroPrep kit (Zymo Research, CA); 80 or 100 ng of RNA were reverse transcribed as described above, and 5 ng of cDNA was analyzed with isoform-specific dPCR assays and QIAcuity Probe PCR Kit according to manufacturer’s instructions (QIAGEN) in either 26k 24-well or 8.5k 96-well Nanoplates (QIAGEN). The plates were analyzed in a QIAcuity digital PCR instrument, using the following PCR program: enzyme activation at 95°C for 2 minutes (1 cycle), denaturation at 95°C for 15 seconds and annealing/extension at 60°C for 30 seconds (40 cycles). The number of different isoforms was quantified by image acquisition of wells according to the selected detection channels in the experiment setup.

### RNA analysis in ROs

Retinal markers *CRX*, *OPN1MW* and *RHO* were used for quality control of ROs; the thresholds were set at >1000 copies/ng RNA. The samples that were below 2 out of 3 thresholds were excluded from the analysis. The thresholds for separation of positive and negative partitions were manually set in all experiments. To correct for different cDNA input, the three identified *ABCA4* isoforms were normalized to the geometric mean of *CRX, RHO and OPN1MW* and the percentage of correct *ABCA4* transcript was calculated as relative to total *ABCA4* within each sample or as relative to total *ABCA4* in wild-type untreated organoids when these were included in the experiment:

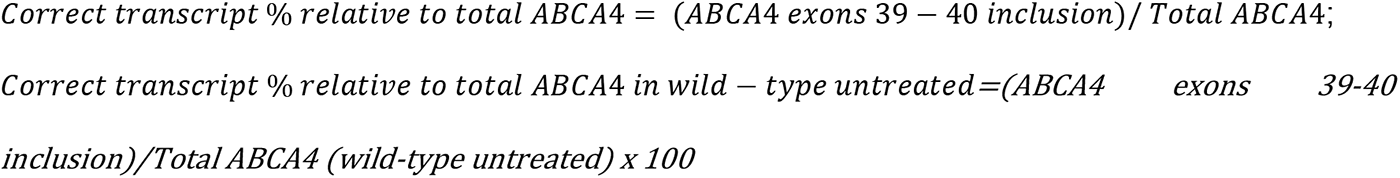

The mean percentage of the detected full-length *ABCA4* isoforms was statistically analyzed using GraphPad Prism 9, with ordinary one-way ANOVA test followed by Dunnet’s multiple comparison test.

To compare the differences in expression of *RHO*, *OPNMW1*, *CRX* and *ABCA4* between batches differentiated with different protocols, the mean expression of each marker was statistically analyzed between each batch with ordinary one-way ANOVA test followed by Tukey’s multiple comparison test.

### Immunohistochemistry (IHC)

The AON-treated ROs were fixed in 2% paraformaldehyde (ThermoFisher Scientific) and 5% sucrose (ThermoFisher Scientific) for 15 minutes at 4°C, followed by a 30-minute incubation in 7.5% sucrose, 30-minute in 15% sucrose and 2-hour incubation in 30% sucrose. The organoids were transferred to a cryomold and embedded in 7.5% gelatin (Porcine skin, Sigma) and 10 % sucrose. The sample blocks were then frozen at -80°C. Sections of 10 µm thick were sliced on a Cryotome FSE (ThermoFisher Scientific), rehydrated in PBS and stained following the protocol described by Cowan et.al [46]. ABCA4 was detected using the anti-ABCA4 3F4 clone (Abcam, 1:100), rhodopsin was stained using the anti-rhodopsin 4D2 clone (Invitrogen, 1:300), mitochondria were detected with an anti-MTCO2 antibody (Abcam, 1:150) and nuclei were stained with Hoechst 33342 (1:1000). Images were collected on an LSM 800 confocal microscope (Carl Zeiss, Germany) using a 60 × objective and analyzed with ZEN Blue edition (Carl Zeiss) using the maximum intensity projection (MIP).

### Identification of protein rescue by Western Blotting

The ROs were pooled (n=10) and lysed in radioimmunoprecipitation assay (RIPA) protein lysis buffer (Abcam, UK) with protease inhibitor cocktail (Roche, Switzerland) and homogenized using a 25-gauge needle. The protein concentration was assessed using the Pierce™ BCA Protein Assay Kit (ThermoFisher Scientific) according to the manufacturer’s instructions and the plate absorbance was read in the SpectraMAX plate reader (Molecular Devices) at 562 nm. The samples (22.5 µg – 85 µg) were loaded on 4-20% Mini-PROTEAN TGX Precast Protein Gels (Bio-Rad), ran for the first 30 minutes at 70 V and the next 4 hours at 100 V in Tris-Glycine SDS buffer. The gels were transferred to PVDF membranes (Millipore) previously activated with methanol, in 1x Tris-Glycine buffer and 20% methanol at 70 mV overnight at 4°C. The membranes were rinsed in PBS-0.1% Tween, blocked in Pure Odyssey Blocking Buffer (Li-COR Biosciences, Lincoln) for 2 hours and incubated with an anti-ABCA4 clone 5B4 (1:1000; Sigma-Aldricht) and anti-Vinculin (1:5000; Abcam) at 4°C overnight. The membranes were washed with PBS-0.1% Tween and incubated with Goat Anti-Mouse IRDye 800 and Goat Anti-Rabbit IRDye 680 (1:5000; Li-COR Biosciences) for 1.5 hours in the dark. The membranes were washed with PBS-0.1% Tween and scanned wet in the Odyssey IR system (Li-COR Biosciences). The intensity of the detected bands was quantified using FIJI ImageJ 1.53c, and the samples were normalized to the wild-type sample. The mean percentage of the detected ABCA4 protein was statistically analyzed using GraphPad Prism 9, with ordinary one-way ANOVA test followed by Dunnet’s multiple comparison test.

## ACKNOWLEDGEMENTS

We are grateful to the patient for donating the cells and participating in this study. We would like to acknowledge the Radboud University Stem Cell facility and Alex Garanto (Radboudumc) for providing the patient-derived IPSCs. We would like to acknowledge Erwin van Wyk (Radboudumc) for providing the pCI-neo.Rho.USH2A-PE40-wt construct and Maarten Holkers (ProQR Therapeutics) for constructing the *ABCA4* minigenes. We thank Carola van Berkel and Maaike van Berkel for supplying gene-edited retinal organoids. We are grateful to Frans Cremers (Radboudumc) for editing the manuscript. This study was financially supported by fundings from Fight for Sight (Grant 5053-5054), European Union’s Horizon 2020 research and innovation programme Marie Sklodowska-Curie Innovative Training Networks (ITN) under grant No. 813490 (StarT), Moorfields Eye Charity, the Wellcome Trust, and the NIHR Biomedical Research Centre based at Moorfields Eye Hospital NHS Foundation Trust and UCL Institute of Ophthalmology.

## AUTHOR CONTRIBUTIONS

Conceptualization: M.K., K.D., R.W.J.C., M.E.C, G.P. and J.S.

Supervision: K.D., R.W.J.C., M.E.C, G.P. and J.S.

Writing-original draft: M.K., P.d.B., T.H.

Methodology: M.K., P.d.B., K.D., T.H., W.B., R.W.J.C., M.E.C. and J.S.

Investigation and data curation: M.K., P.d.B., D.P., S.E.L, K.D., T.H., R.W.J.C., M.E.C. and J.S.

Funding acquisition: A.R.W., R.W.J.C., M.E.C. and G.P.

Writing-review & editing: M.K., P.d.B., D.P., T.H., W.B., R.W.J.C., M.E.C. and J.S.

## FIGURE LEGENDS SUPPLEMENTARY DATA

**Figure S1.**
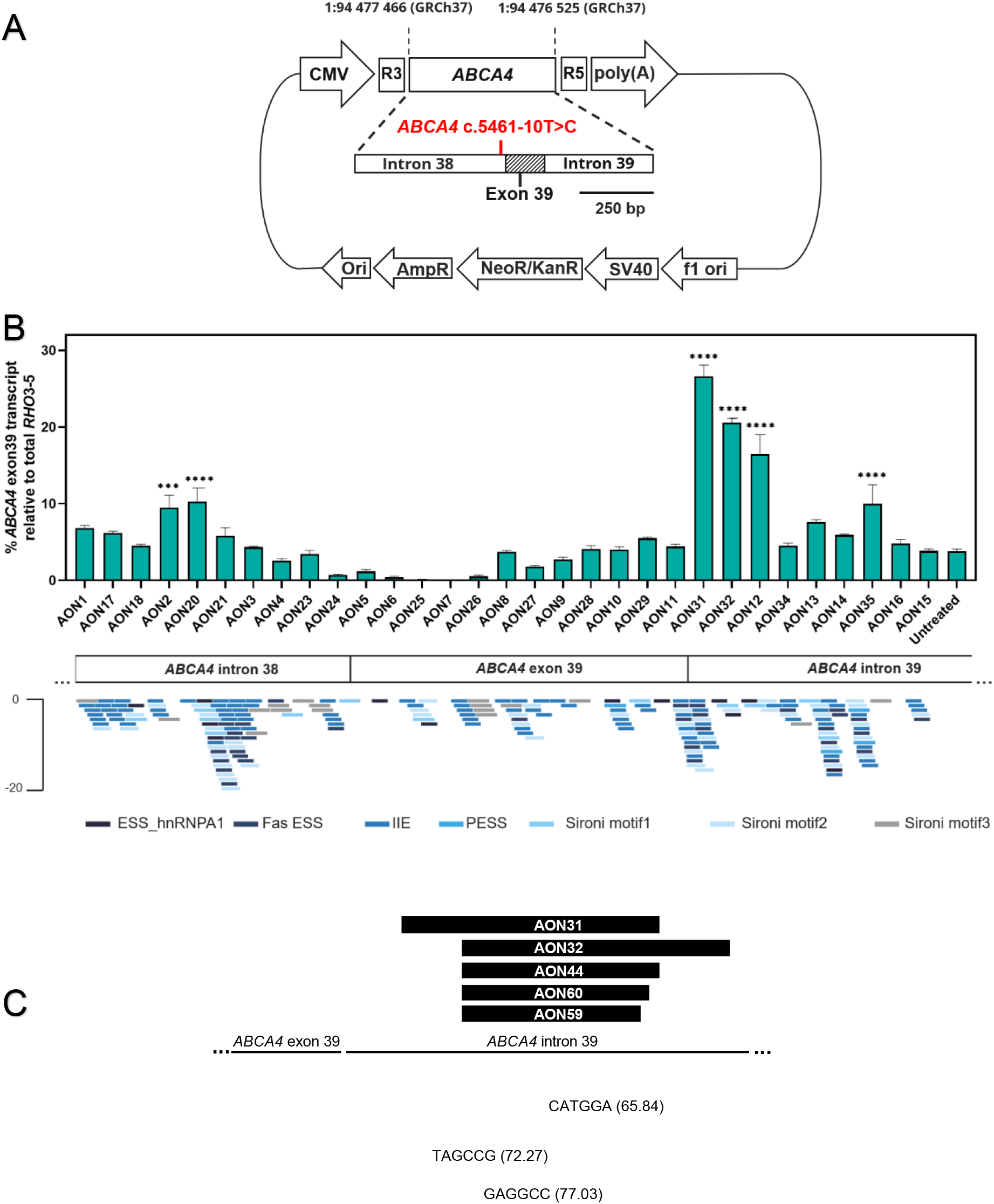
Splicing correction in minigene-transfected HEK293T cells treated with AONs targeting the intronic splicing silencers to correct the aberrant splicing caused by *ABCA4* c.5461-10T>C. (A) Schematic representation of the minigene construct showing the *ABCA4* exon 39 flanked by parts of adjacent introns; the rest of the plasmid backbone was the same as shown in Figure 2A. (B) The first AON screen contained 31 different AONs with the 2’O-Methyl (2’OMe) chemistry and phosphorothioate (PS) backbone; the graph displays the percentages of splicing rescue after 100nM AON transfection treatment on cells transfected with 50 ng of minigene (n=2). The transcript analysis, assessed 24 hours post-treatment, suggested that the AONs targeting the intron 39 region demonstrated the most significant therapeutic effect on the splicing modulation. AON31 and AON32 showed the most potent splicing rescue by increasing the levels of wild-type transcript up to 26% and 20%, respectively; therefore, these two AONs served as model molecules for design of optimized AONs. Below are displayed the binding sites for RNA-splicing proteins obtained fromHuman Splicing Finder (Desmet et al. 2009). ***p≤0.001, ****p≤0.0001, ordinary one-way ANOVA test followed by Dunnet’s multiple comparison test. The details of each binding site can be found in Table S2. (C) Lead candidates AON 31 and AON32 and their shorter versions AON44, AON60 and AON59 are complementary to the intron 39 region where three strong splicing silencer motifs are located that are involved in the recruitment of the splicing protein heterogeneous nuclear ribonucleoprotein A1 (hnRNP A1). The AONs block the motifs and thereby enhances the splicing in favor of exon 39 and 40 re-inclusion.

**Figure S2.**
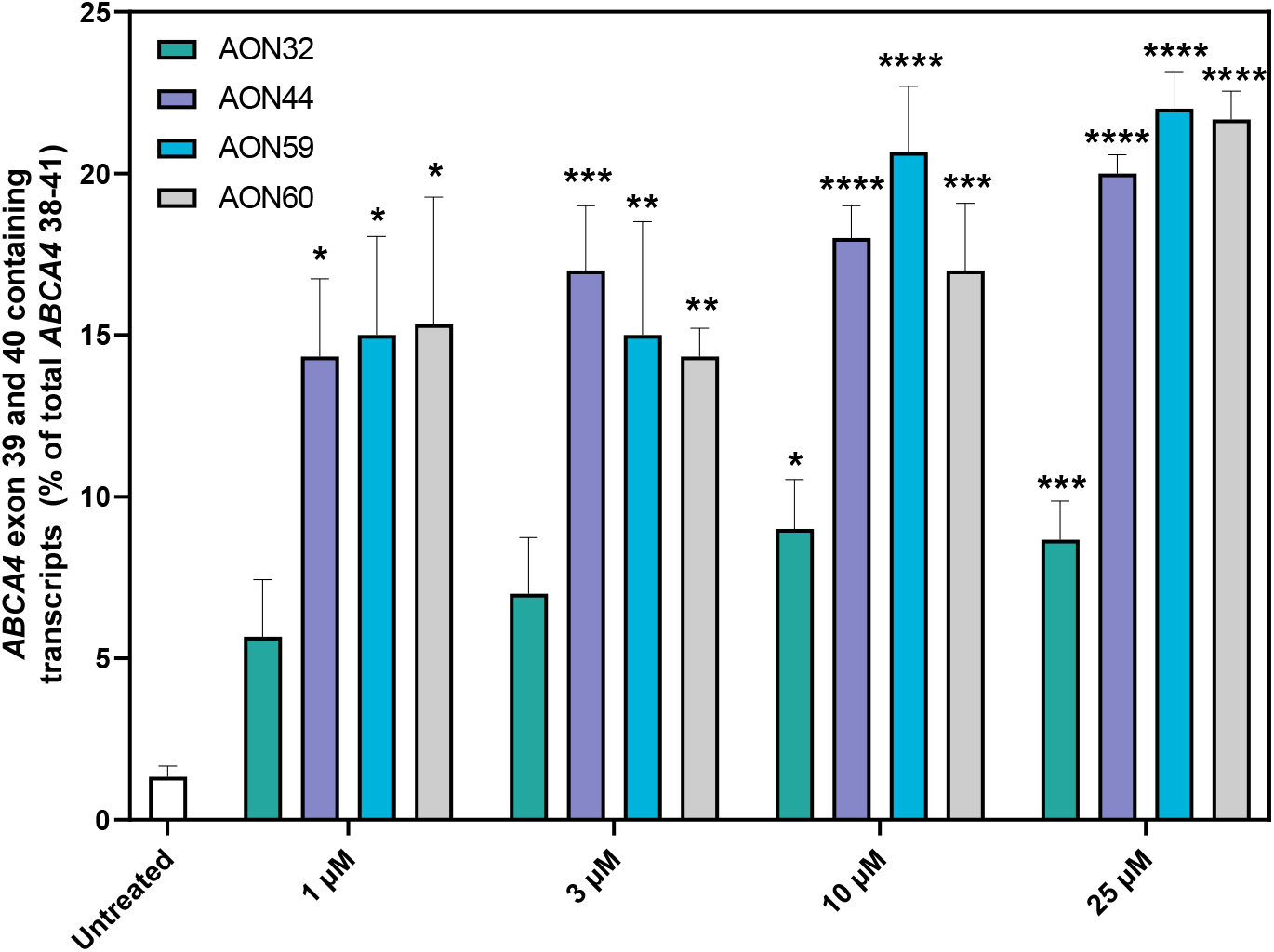
The dose-response curve in midigene-transfected. (Figure 2A) HEK293 cells treated gymnotically with 1, 3, 10 or 25 µM concentration for X hours of best selected AONs: AON44, AON59 and AON60. AON32 served as control since it consists in the long version of other selected AONs. A clear concentration-dependent effect of all used AONs is observed. Data are shown as mean±s.e.m., n=3. All samples were compared to the untreated sample, *p≤0.05, **p≤0.01, ***p<0.001, ****p<0.0001, ordinary one-way ANOVA test followed by Dunnet’s multiple comparison test.

**Figure S3.**
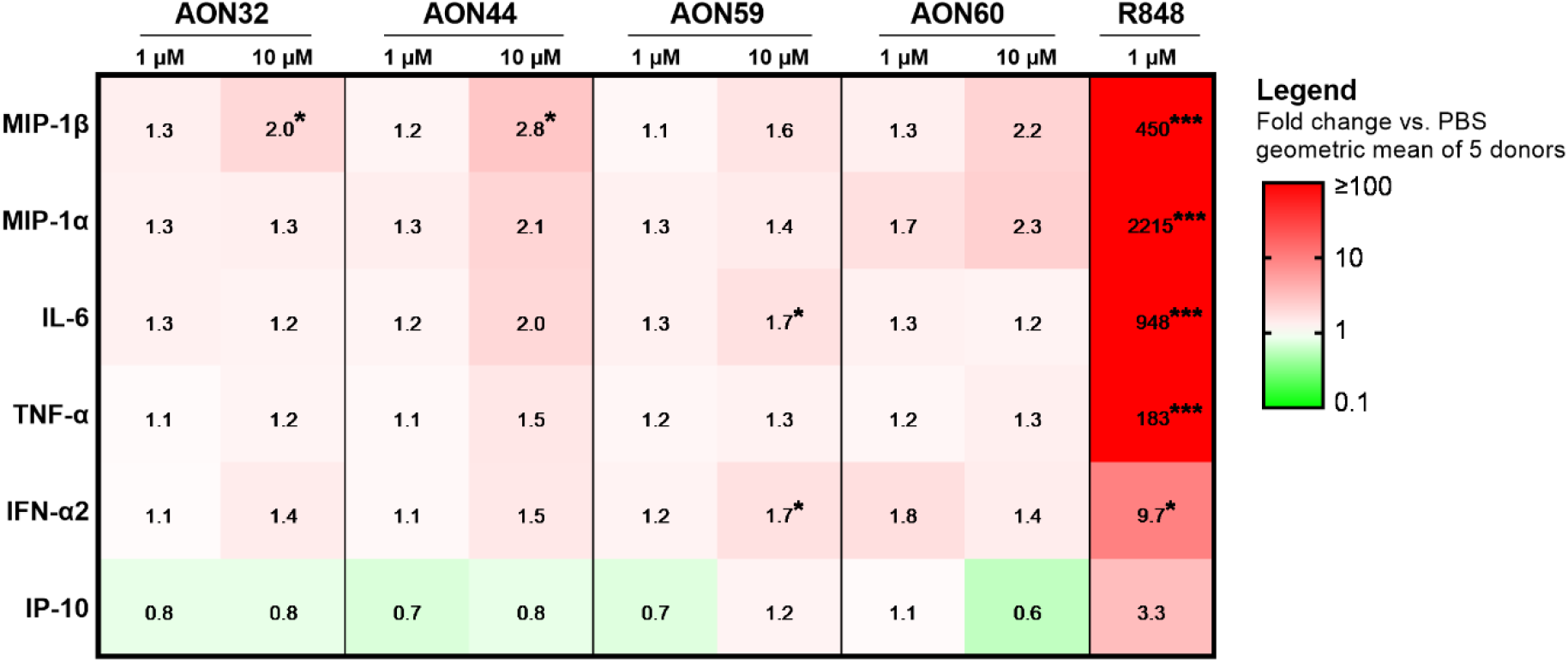
Immunostimulatory potential of lead AON candidates. The heatmap depicts the fold change levels of cytokine concentrations in culture supernatant after 48-hour exposure to lead AON candidates or positive control R848 as compared to PBS-treated human peripheral blood mononuclear cells. Cell fill colors indicate the direction and the degree of the fold change. Positive control R848 resulted in significantly increased concentrations of all measured cytokines except for IP-10. All AONs were shown to exert a slight influence on cytokine secretion, reaching statistical significance for AON32 [10 µM], AON44 [10 µM] (MIP-1β) and AON59 [10 µM] (IL-6 and IFN-α2). Statistical testing was performed on log-transformed concentration values using mixed-model analysis, applying Dunnett’s correction for multiplicity. Statistically significant differences vs. PBS were annotated with *p<0.05, **p<0.01, ***p<0.001.

**Figure S4.**
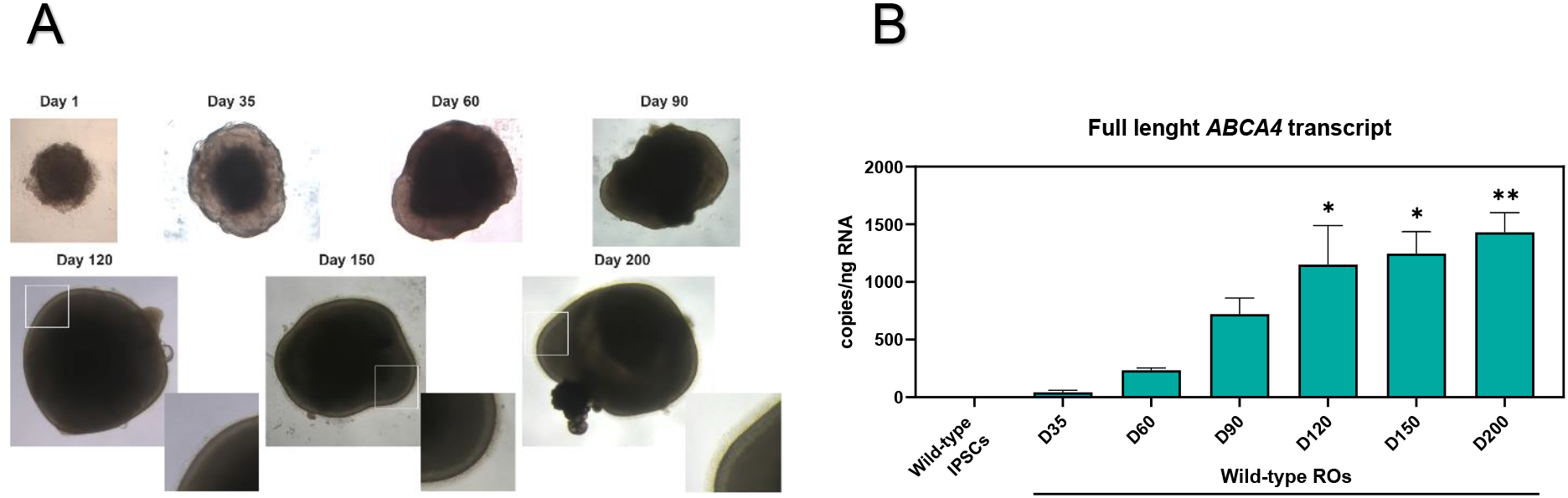
Morphological and ABCA4 transcript analysis of wild-type ROs over time. (A) Control wild-type ROs were generated from wild-type iPSCs that served as parent isogenic line for generation of gene edited homozygous *ABCA4* c.5461-10T>C iPSCs. The cells display a round clump already at day 1 that develops in an organoid with dark core and sharp edges (day 35). The development of the neural retina was observed at day 90, and by day 120, the organoids were surrounded by a brush border that contains the inner and outer segments of photoreceptor cells. (B) The isoform analysis detected *ABCA4* isoforms only 35 days after organoid differentiation; the *ABCA4* expression increases at day 60 and 90 to 120, after which the expression is significantly higher over the expression detected in IPSCs. Moreover, 120 days after differentiation, the expression of *ABCA4* remained relatively constant. Data are shown as mean±s.e.m., n=6. Statistically significant differences vs. wild-type IPSCs were annotated with *p≤0.05, **p≤0.01, ordinary one-way ANOVA test followed by Dunnet’s multiple comparison test.

**Figure S5.**
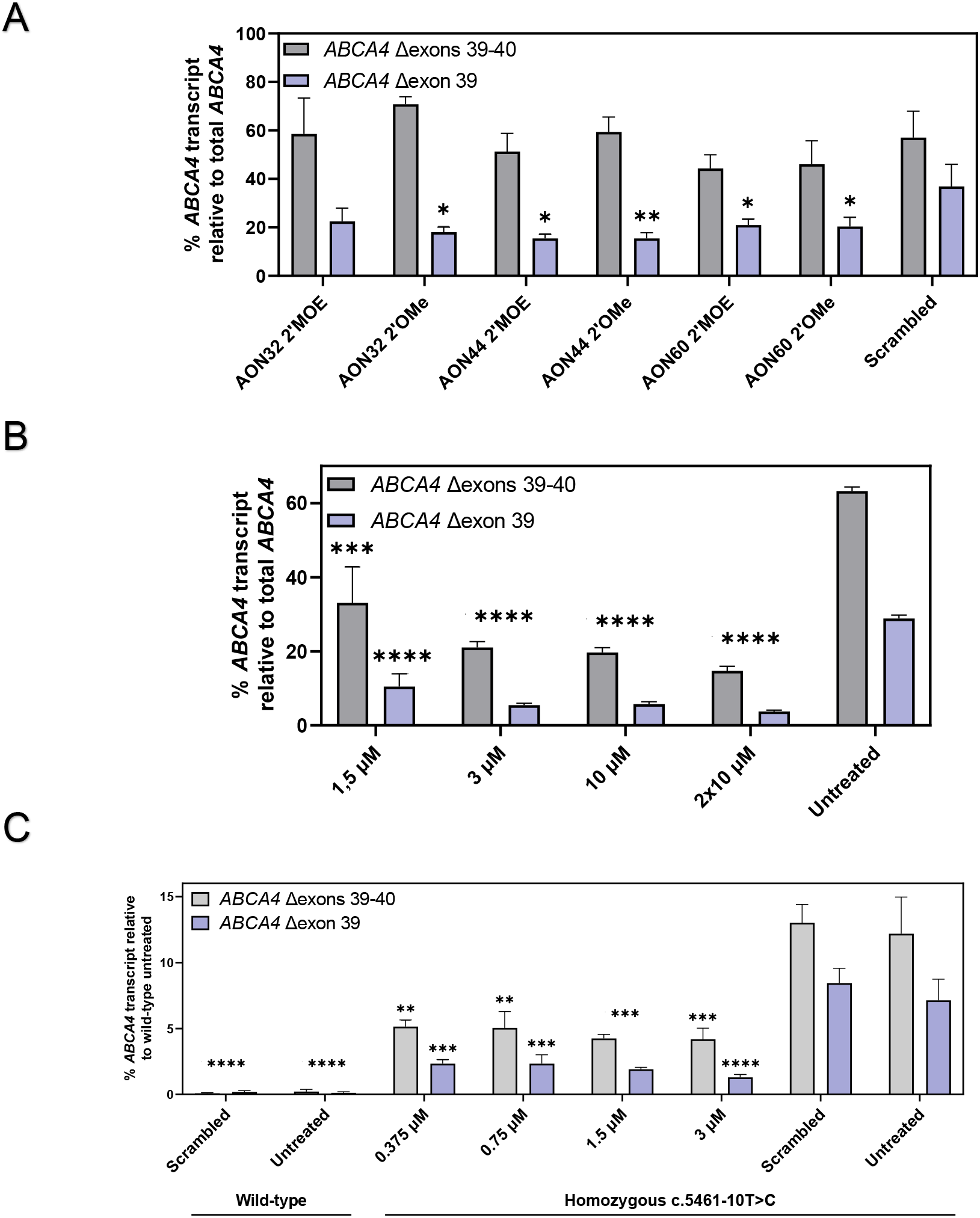
Overview of the percentage of truncated *ABCA4* isoforms in organoid studies. (A) Percentages of single skip and double skip *ABCA4* isoforms in the organoid study described in Figure 4A, (B) Figure 4B and (C) Figure 6A. All samples bi-allelic for *ABCA4* c.5461-10T>C showed considerably higher amounts of *ABCA4* isoforms with double exon skip than single exon skip. On the other hand, the AON-treatment seems to exhibit restoration of both truncated isoforms. Data are shown as mean ± s.e.m. Statistically significant differences vs. untreated or scrambled were reported as *p≤0.05, **p≤0.01, ***p<0.001, ****p<0.0001, ordinary one-way ANOVA test followed by Dunnet’s multiple comparison test. n=6 per condition.

**Figure S6.**
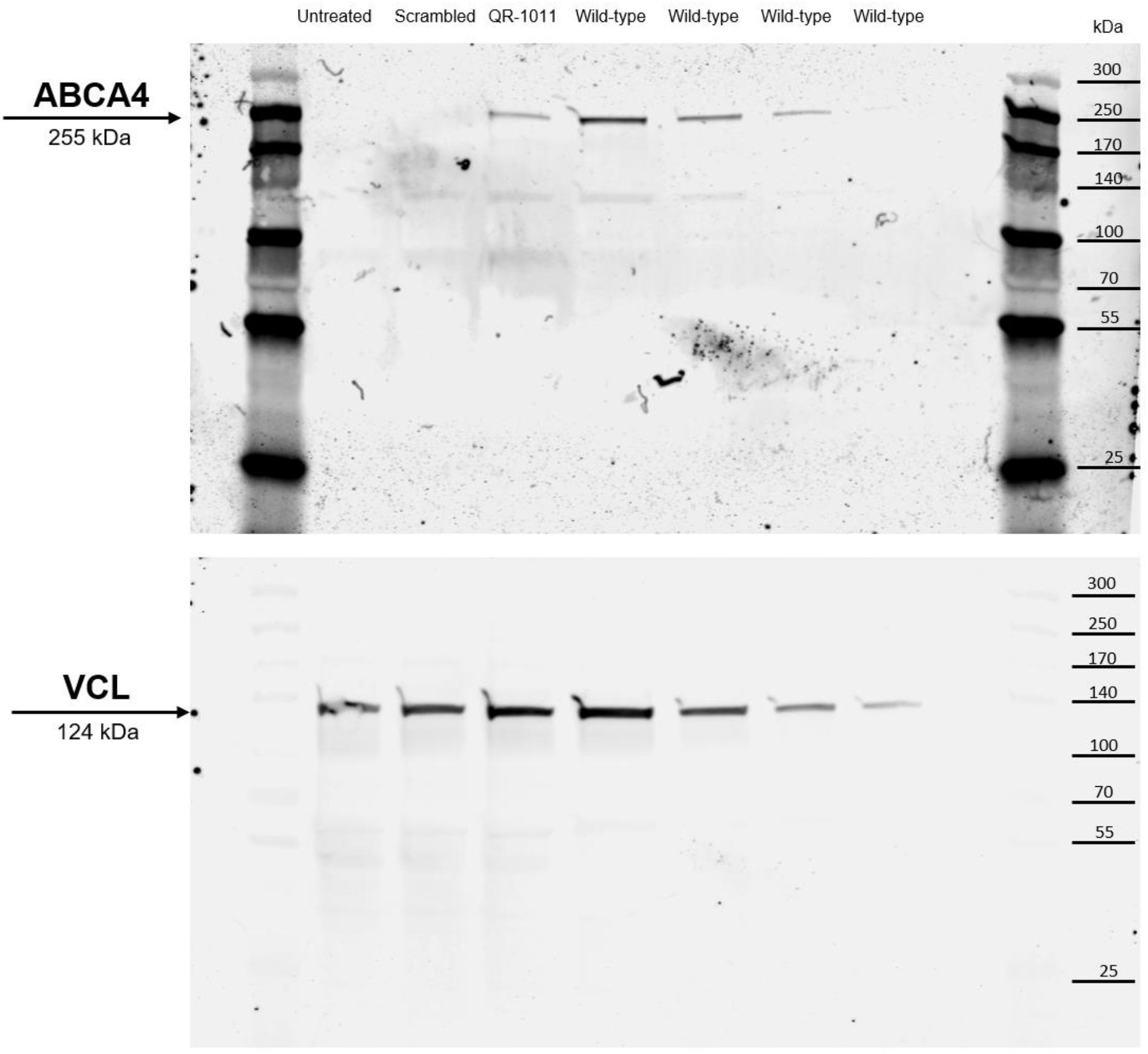
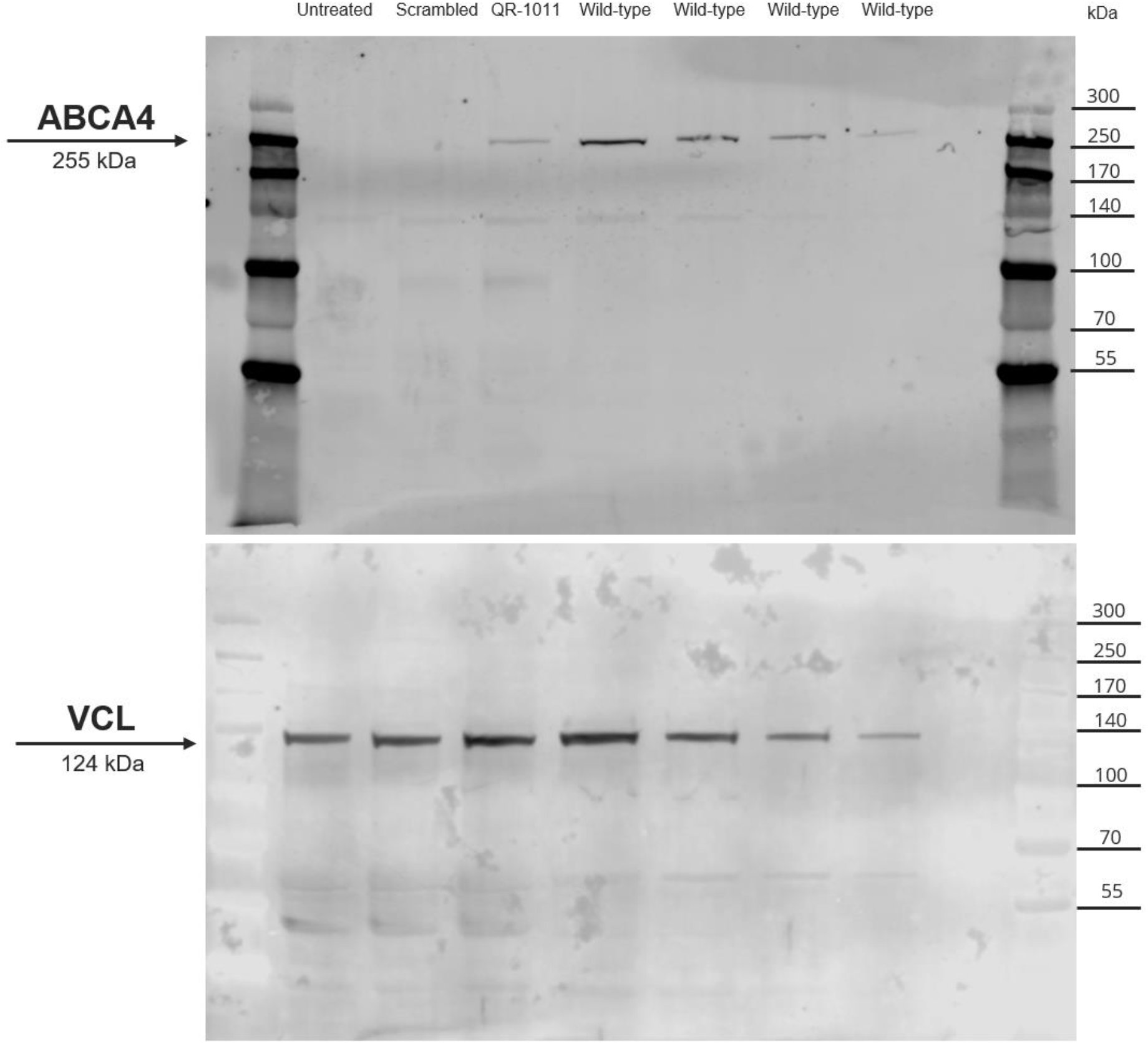
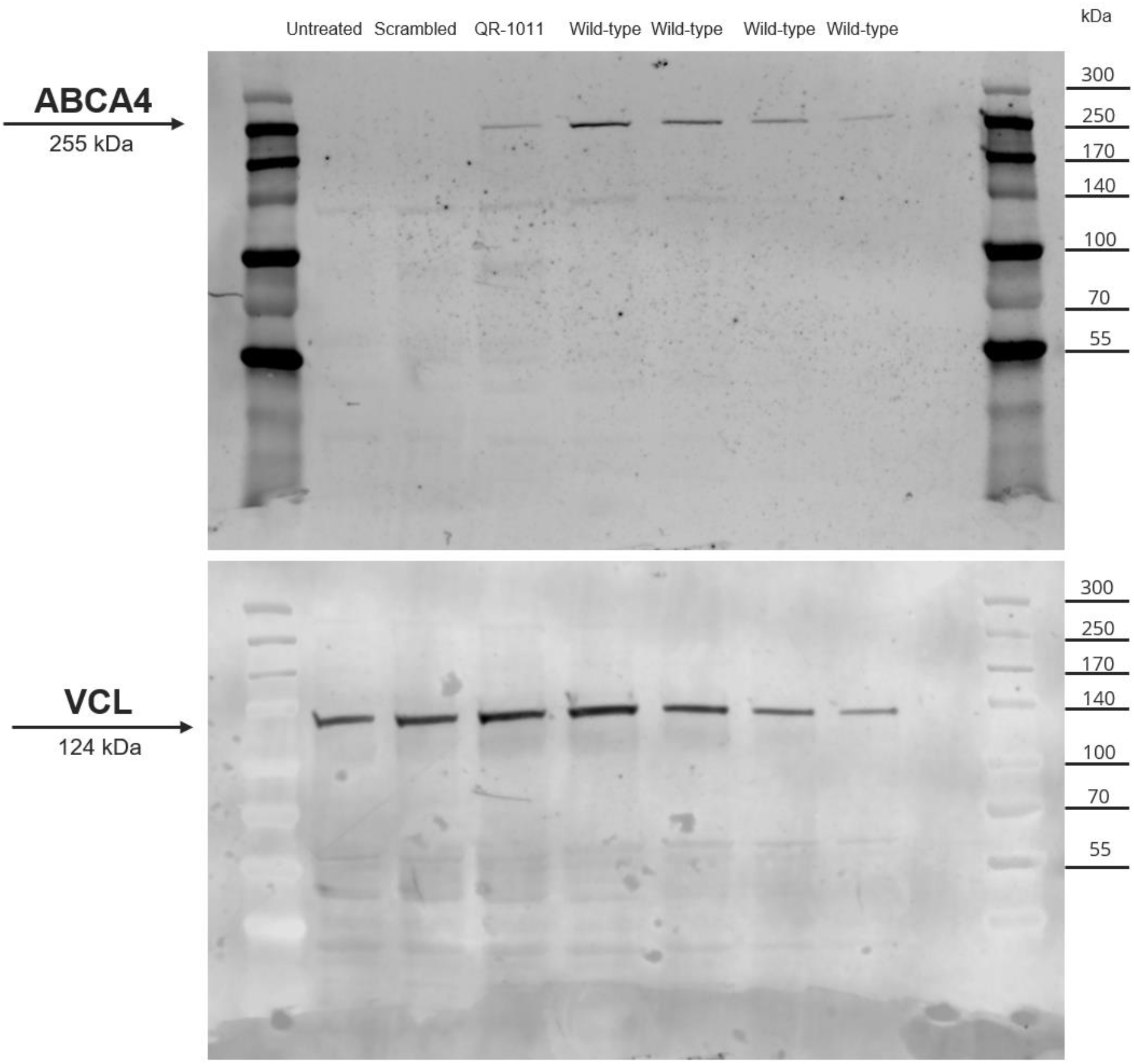
Technical replicates of Western Blots with protein lysates from patient-derived ROs homozygous for -10T>C and wild-type ROs. Wild-type ABCA4 protein was regenerated in patient-derived ROs after a 3 µM dose of QR-1011 in an 8-week long treatment. The three technical replicates suggest that no wild-type ABCA4 protein was present in untreated STGD1 organoids and those treated with the scrambled oligo (n=10 per condition).

